## Supplementary Figures for "A statistical framework for assessing pharmacological response and biomarkers using uncertainty estimates"

### GDSC Screening Team: Howard Lightfoot, Wanjuan Yang, Maryam Soleimani, Syd Barthorpe, Tatiana Mironenko, Alexandra Beck, Laura Richardson, Ermira Lleshi, James Hall, Charlotte Tolley, William Barendt

### Supplementary Figures

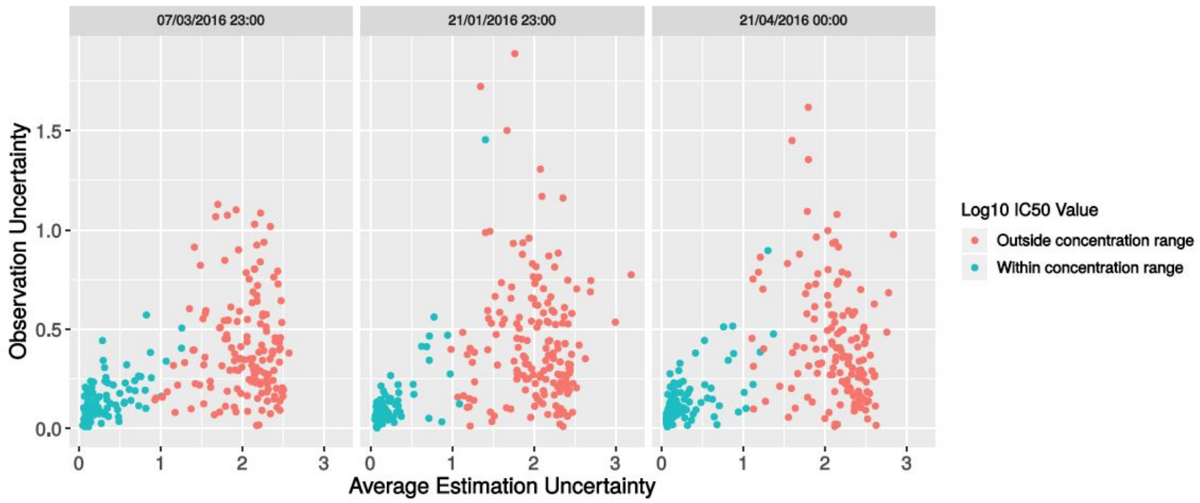


**Supplementary Figure S1: Investigation of batch effects in the replicate data.** Scatterplot of observation uncertainty against average estimation uncertainty, split by experimental batch.

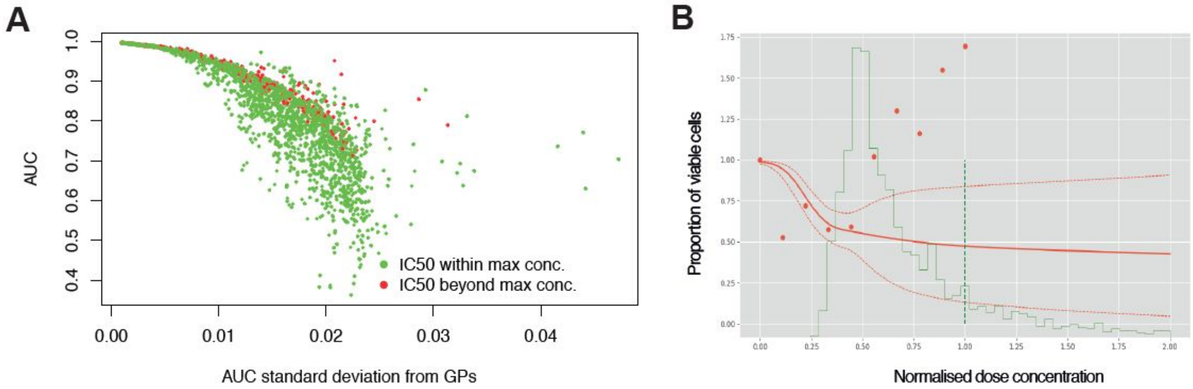


**Supplementary Figure S2: High estimation uncertainty independent of concentration range. (A)** Estimation uncertainty for AUC (standard deviation) across replicate experiments were robust to whether the $IC_{50}$ was within or beyond the maximum concentration tested. **(B)** Dose response of a single experiment where there was high estimation uncertainty (dotted red line) despite the fitted curve (red line) crossing 50% viable cells before reaching the max dose concentration (dotted green line). The probability distribution of $IC_{50}$ is estimated across dose concentrations (solid green line), however, raw data points (red dots) show increased cell viability with increased dose.

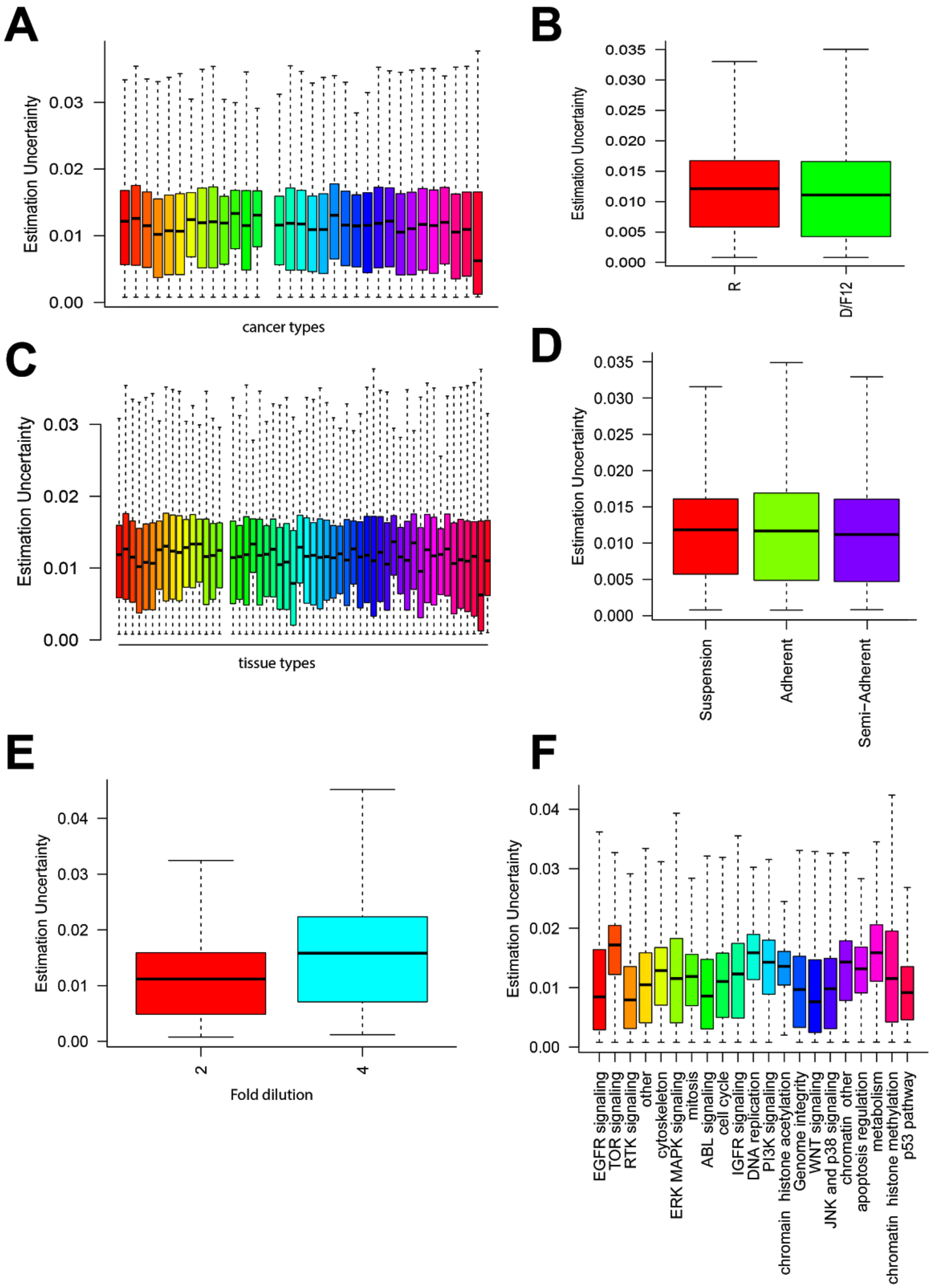


**Supplementary Figure S3: AUC estimation uncertainty grouped by (A)** cancer type of cell lines, **(B)** growth media used, **(C)** tissue of origin, **(D)** growth condition of the cells, **(E)** dilution factor for each dose tested of a drug, and **(F)** target pathways of drugs.

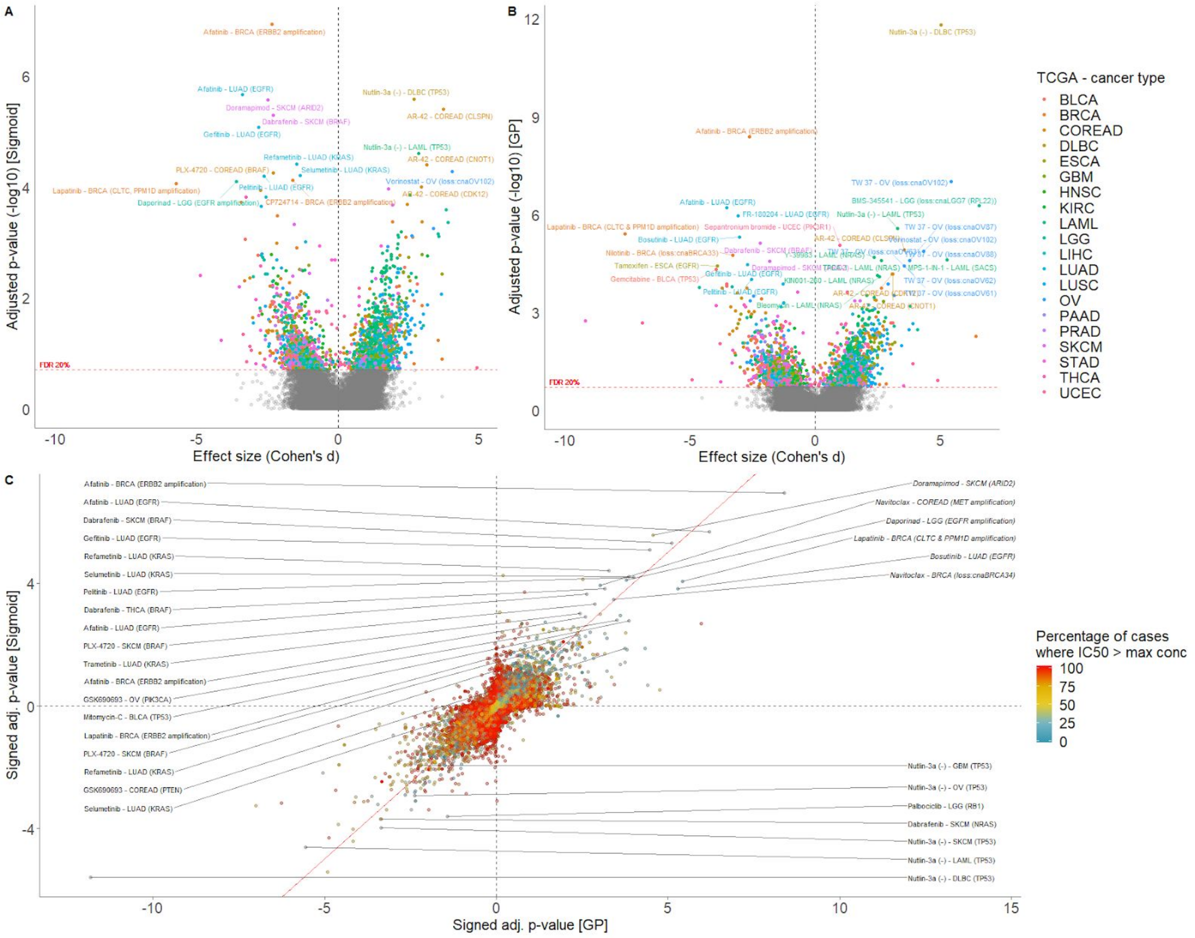


**Supplementary Figure S4: Comparison of sigmoid and GP curve fitting.** Volcano plot of drug response biomarker associations based on **(A)** sigmoid and **(B)** GP curve fitting. **(C)** Drug response biomarker comparison based on both curve fittings, and color coding percentage of drug response data observed within concentration range.

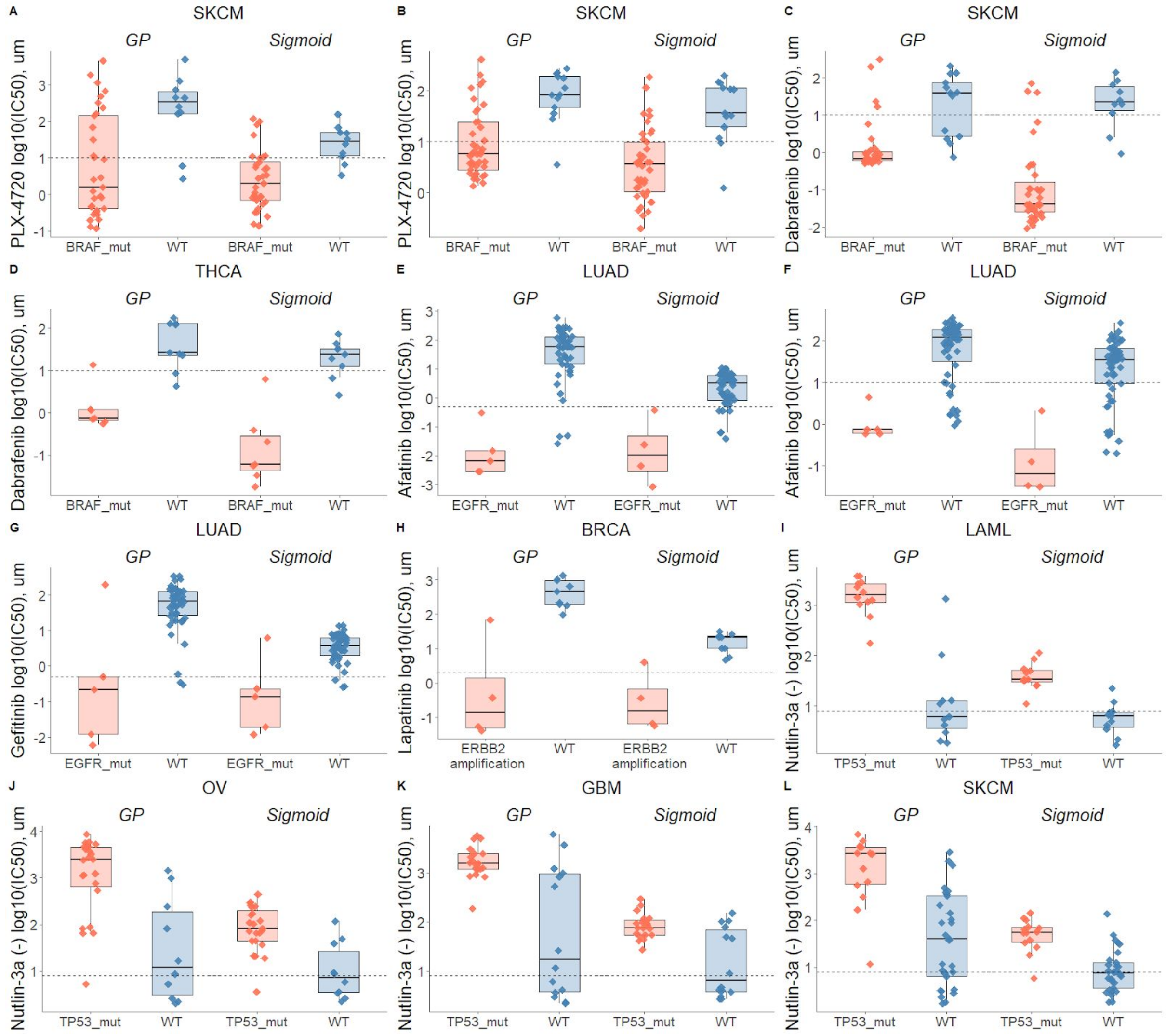


**Supplementary Figure S5: Drug response biomarker comparison based on both curve fittings.** Dashed line depicts maximum concentration. **(A)** Skin cutaneous melanoma (SKCM) treated with PLX-4720 (BRAF inhibitor) and stratified with *BRAF* mutations; **(B)** Skin cutaneous melanoma (SKCM) treated with PLX-4720 (BRAF inhibitor) and stratified with *BRAF* mutations - replicate; **(C)** Skin cutaneous melanoma (SKCM) treated with Dabrafenib (BRAF inhibitor) and stratified with *BRAF* mutations; **(D)** Thyroid carcinoma (THCA) treated with Dabrafenib (BRAF inhibitor) and stratified with *BRAF* mutations; **(E)** Lung adenocarcinoma (LUAD) treated with Afatinib (ERBB2, EGFR inhibitor) and stratified with *EGFR* mutations; **(F)** Lung adenocarcinoma (LUAD) treated with Afatinib (ERBB2, EGFR inhibitor) and stratified with *EGFR* mutations - replicate; **(G)** Lung adenocarcinoma (LUAD) treated with Gefitinib (EGFR inhibitor) and stratified with *EGFR* mutations; **(H)** Breast invasive carcinoma (BRCA) treated with Lapatinib (ERBB2, EGFR inhibitor) and stratified with *ERBB2* amplifications; **(I)** Acute Myeloid Leukemia (LAML) treated with nutlin-3a (MDM2 inhibitor) and stratified by *TP53* mutants; **(J)** Ovarian serous cystadenocarcinoma (OV) treated with nutlin-3a (MDM2 inhibitor) and stratified by *TP53* mutants; **(K)** Glioblastoma multiforme (GBM) treated with nutlin-3a (MDM2 inhibitor) and stratified by *TP53* mutants; **(L)** Skin cutaneous melanoma (SKCM) treated with nutlin-3a (MDM2 inhibitor) and stratified by *TP53* mutants.

#### Supplementary Tables

1. Supplementary Table S1 - Summary of pharmacogenomic associations based on ANOVA
2. Supplementary Table S2 - Pharmacogenomic associations based on Bayesian testing of GP curve fits
3. Supplementary Table S3 - Raw and curve fitted replicate dataset
